# The CUT&RUN Blacklist of Problematic Regions of the Genome

**DOI:** 10.1101/2022.11.11.516118

**Authors:** Anna Nordin, Gianluca Zambanini, Pierfrancesco Pagella, Claudio Cantù

## Abstract

Cleavage Under Targets and Release Using Nuclease (CUT&RUN) is an increasingly popular technique to map genome-wide binding profiles of histone modifications, transcription factors and co-factors. The ENCODE project and others have compiled blacklists for ChIP-seq which have been widely adopted: these lists contain regions of high and unstructured signal, regardless of cell type or protein target. While CUT&RUN obtains similar results to ChIP-seq, its biochemistry and subsequent data analyses are different. We found that this results in a CUT&RUN-specific set of undesired high-signal regions. For this reason, we have compiled blacklists based on CUT&RUN data for the human and mouse genomes, identifying regions consistently called as peaks in negative controls by the CUT&RUN peak caller SEACR. Using published CUT&RUN data from our and other labs, we show that the CUT&RUN blacklist regions can persist even when peak calling is performed with SEACR against a negative control, and after ENCODE blacklist removal. Moreover, we experimentally validated the CUT&RUN Blacklists by performing reiterative negative control experiments in which no specific protein is targeted, showing that they capture >80% of the peaks identified. We propose that removing these problematic regions prior to peak calling can substantially improve the performance of SEACR-based peak calling in CUT&RUN experiments, resulting in more reliable peak datasets.

## Introduction

Cleavage Under Targets and Release Using Nuclease (CUT&RUN, hereafter referred to as C&R) is a technique developed to map the genome-wide binding profiles of histone modifications, transcription factors and co-factors [1]. Like other genomics techniques, C&R uses short-read next generation sequencing (NGS) to generate datasets, and thus depends on reliable mapping and genome assemblies for their accurate interpretation. Repetitive regions, assembly gaps, and other computational challenges can lead to the rise of sequencing artifacts, resulting in areas of significant signal enrichment that are not due to biological processes [2]. Additionally, high-signal regions can occur as consequence of several procedures, such as cell fixation or PCR-based amplification after adaptor ligation of sequencing libraries [3–5].

Sets of artifactual and high-signal regions that should be excluded from the analysis, known as blacklists, have previously been generated by the ENCODE consortium and others for chromatin immunoprecipitation followed by sequencing (ChIP-seq) [6, 7]. C&R and ChIP-seq obtain comparable results, and the ENCODE blacklist have been used for both these assays. However, C&R and ChIP rely on different biochemical procedures; for example, in ChIP-seq the chromatin is crosslinked, sonicated, and immunoprecipitated, while in C&R no crosslinking is applied and the target-associated DNA fragments are obtained by digestion with the MNase-proteinA fusion protein [1]. The determination of signal enrichment, peak calling, can also be done differently between the two techniques. While most ChIP-seq peak calling algorithms use local signal to background ratios to determine significance, the low background typical of C&R renders this difficult. This led to the development of the C&R specific peak caller SEACR, which uses global background (as opposed to local background) to determine a signal to noise threshold against which to call peaks [8]. Artifacts in ChIP-seq have been shown to bias peak calling results [2]; we reasoned that, due to its reliance on global background signal, SEACR may be even more sensitive to unspecific signals.

To address these differences between C&R and other NGS-based techniques, we compiled blacklists for the hg38 human and mm10 mouse genomes built exclusively on C&R data. To determine regions that produce detectable signals in negative controls, hence likely to be erroneously called as peaks across experiments, we downloaded C&R negative control datasets (N=20 per genome, from 40 different studies), and performed peak calling on them to identify peaks consistently identified by SEACR. This allowed us to establish a human and a mouse C&R blacklist of by-definition artifactual peaks. Both human and mouse C&R blacklists obtained in this manner show high precision, as they encompass less than 0.2% of their respective genome yet succeed – when used to subtract the false peaks from C&R datasets – in increasing the genome-wide variance between samples, indicating that the blacklist regions are commonly enriched across C&R experiments, regardless of the target and the cell type. This indicates that these regions ought to be removed.

## Results and Discussion

### Generation of the Blacklists

We compiled human and mouse genomic C&R blacklists by downloading publicly available raw C&R data from the European Nucleotide Archive (https://www.ebi.ac.uk/ena/browser/home) and processing them to identify high signal regions (Figure 1A). We included 40 negative controls (20 for human and 20 for mouse) from 40 different studies to ensure diversity on the model systems used and identify cell-type agnostic regions (Materials and Methods Table 1). The criteria for inclusion were a clear labelling system as C&R negative controls (i.e., “no antibody”, “IgG”, etc.), and the use of pair-end NGS technology. We performed trimming on the data to remove adapters and then mapped to the human (hg38) or mouse (mm10) genomes with bowtie2, using commonly recommended settings for C&R data [9, 10]. Once mapped, we deduplicated the BAM files to stringently identify artifacts that occur regardless of PCR duplication rates. We applied peak calling of these negative control samples with SEACR on stringent mode, using a threshold of 0.001 to identify the highest 0.1% signals. We then extended the peak regions by 1000 bp in either direction, to ensure capturing peaks that may be slightly shifted between datasets and overlapped the peak sets resulting from each different individual sample to measure the reproducibility of false positive signal. The blacklists are compiled by peaks present in at least 30% of the negative controls used: we consider this probability sufficiently high to indicate an unacceptable chance of false positive hits.

**Figure 1:**
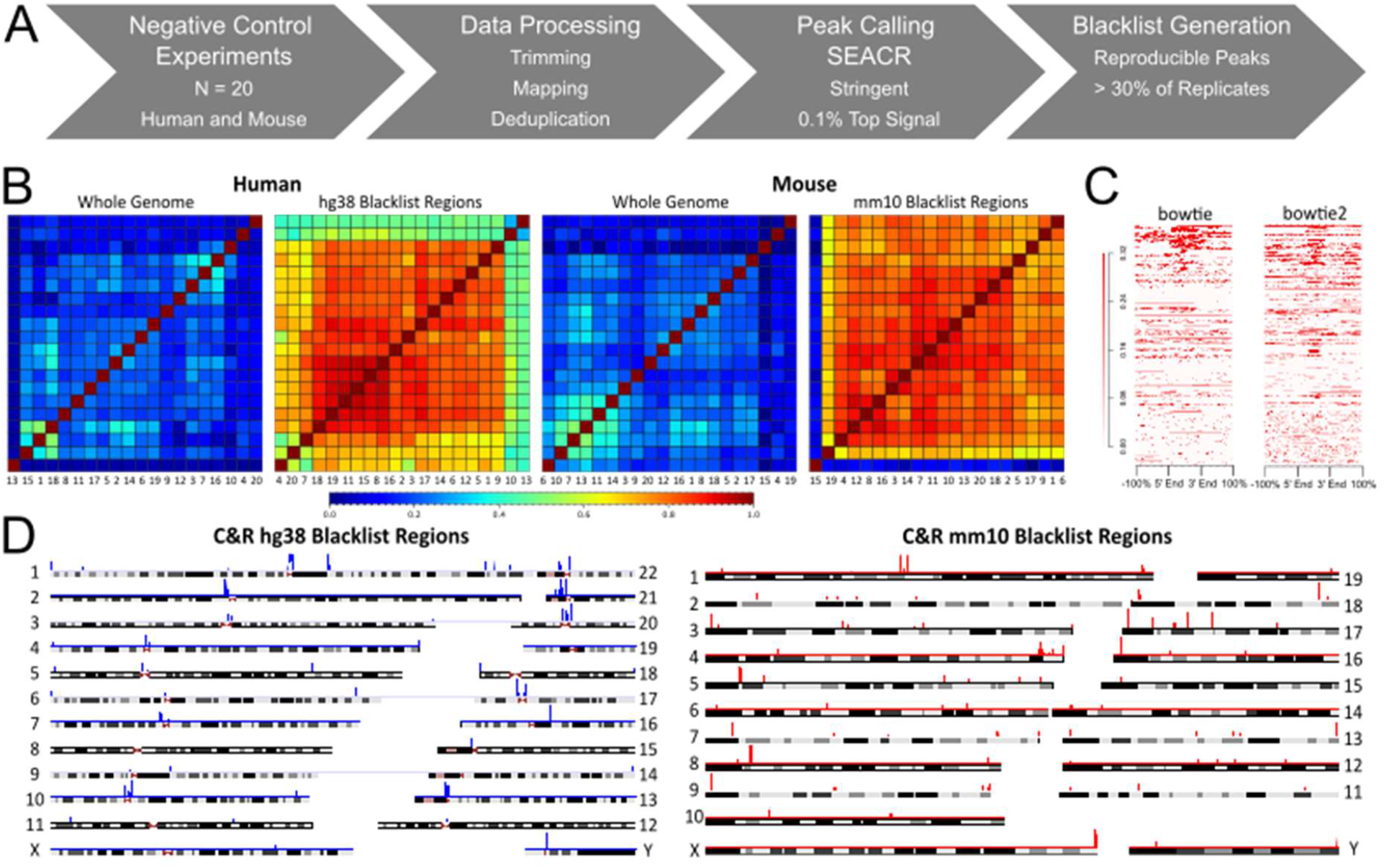
Generation of the CUT&RUN Blacklists. **A.** Schematic of blacklist generation strategy. **B.** Plots of Spearman’s correlation between negative control samples (N=20 each) in human and mouse, showing genome-wide correlation and correlation within the C&R blacklist regions in hg38 and mm10 genomes. In both human and mouse, samples show dramatically increased correlation when considering only the blacklist regions. **C.** Signal enrichment plots of the hg38 blacklist regions for a representative human negative control sample, after mapping with bowtie or bowtie2. More stringent mapping with bowtie did not drastically change the enrichment in the blacklisted regions. **D.** Chromosome maps showing genome wide locations of regions contained in the C&R hg38 and mm10 blacklists.

**Table 1:** Data Availability and References.

| Human |  |  |  | Mouse |  |  |  |
| --- | --- | --- | --- | --- | --- | --- | --- |
| Number | Project | Sample | Reference | Number | Project | Sample | Reference |
| 1 | PRJNA758691 | SRR15663649 | [18] | 1 | PRJNA682243 | SRR13188236 | [19] |
| 2 | PRJNA753322 | SRR15403020 | [20] | 2 | PRJEB41862 | ERR4973502 | [21] |
| 3 | PRJNA699604 | SRR13634070 | [22] | 3 | PRJNA777234 | SRR16694279 | [23] |
| 4 | PRJNA655230 | SRR12387746 | [24] | 4 | PRJNA744774 | SRR15069561 | [25] |
| 5 | PRJNA427801 | SRR6426139 | [26] | 5 | PRJNA719369 | SRR14134905 | GSE171383, |
| 6 | PRJNA776721 | SRR16674836 | [27] | 6 | PRJEB51482 | ERR9130874 | E-MTAB-11513 |
| 7 | PRJNA756549 | SRR15539083 | [28] | 7 | PRJNA746301 | SRR15123732 | [29] |
| 8 | PRJNA721947 | SRR14238385 | [30] | 8 | PRJNA722185 | SRR14243166 | [31] |
| 9 | PRJNA691366 | SRR13586925 | [32] | 9 | PRJNA753786 | SRR15414824 | [33] |
| 10 | PRJNA816825 | SRR18337644 | [34] | 10 | PRJNA744230 | SRR15054366 | [35] |
| 11 | PRJEB55317 | ERR10047705 | [36] | 11 | PRJNA786482 | SRR17139040 | [37] |
| 12 | PRJNA836267 | SRR19139489 | [38] | 12 | PRJNA860380 | SRR20325067 | [39] |
| 13 | PRJNA798000 | SRR17642967 | [40] | 13 | PRJNA656290 | SRR12424468 | [41] |
| 14 | PRJNA717224 | SRR14068376 | [42] | 14 | PRJNA862741 | SRR20665931 | [43] |
| 15 | PRJNA704964 | SRR13785931 | [44] | 15 | PRJNA658977 | SRR12507586 | [45] |
| 16 | PRJNA682426 | SRR13194196 | [46] | 16 | PRJNA678949 | SRR13073031 | [47] |
| 17 | PRJNA888075 | SRR21837879 | [48] | 17 | PRJNA527826 | SRR8745654 | [49] |
| 18 | PRJNA413473 | SRR6144305 | [50] | 18 | PRJNA682340 | SRR13190129 | [51] |
| 19 | PRJNA647352 | SRR12267593 | [52] | 19 | PRJNA864644 | SRR20727958 | [53] |
| 20 | PRJNA562266 | SRR10022374 | [54] | 20 | PRJNA493794 | SRR7939979 | [55] |

Genome-wide, the datasets show very low correlation (Figure 1B), which is expected due to random distribution of MNase-based digestion when a non-targeting antibody is used [9]. However, when considering only the blacklisted regions the correlation is high (Figure 1B), indicating that these regions are enriched across independent negative control samples and ought to be considered as artifactual. To ensure that the blacklists are valid regardless of the data processing pipeline, we applied more stringent mapping with bowtie – instead of bowtie2 – allowing neither sequence mismatches nor multimapping. Though the fraction of reads within the blacklist peaks (FRIP score; a typical measure of the efficiency of the experiment) typically decreased with more stringent mapping (Supplementary Table 1), the average signal enrichment profile across blacklist regions was comparable and still enriched compared to background (Figure 1C), demonstrating that a majority of these problematic regions display real signal enrichment independently from mapping biases.

The blacklisted regions are scattered throughout the genome, both in human and in mouse, and involve all chromosomes (Figure 1D). The C&R blacklists also contain the mitochondrial genome, as well as regions from unknown or random chromosomes which are mappable with bowtie2, as these regions can contribute to noise in experiments.

### Comparison of the C&R Blacklists with the ENCODE Blacklists

When compared with the latest version (v2) of the ENCODE blacklists [6], our C&R blacklists cover a smaller portion of the genome (Figure 2A). For the comparison, we manually added the mitochondrial genome to the ENCODE blacklists, as this is recommended to remove but not included in the list. Several ENCODE blacklist regions are broader than the C&R blacklist regions; the ENCODE blacklisted regions were in fact extended to encompass unmappable or neighboring high-signal regions, and blacklist regions within a certain distance range were merged [6]. In contrast, we chose not to manually curate or merge the identified regions, or extend them beyond 1000 bp in either direction, to provide the dual advantage of i) increasing the specificity in identifying the high-signal C&R false peaks and ii) removing as little of the genome as possible. Therefore, one ENCODE blacklist peak may overlap with multiple C&R blacklist peaks. While due to their different biochemistries ChIP-seq and C&R can produce different sets of false positives, we noticed that several false peaks are shared between the ENCODE and C&R blacklists (Figure 2B). We interpret that these artifacts are caused by genome assembly and mapping biases rather than the undesired purification and identification of unspecific DNA fragments [6]. Moreover, the overlap between the ENCODE and the C&R blacklists constitutes an important validation of our approach. On the other hand, as expected, there are C&R specific regions absent in the ENCODE lists (Figure 2C), and many regions included in the ENCODE blacklists do not show any signal enrichment in the C&R negative controls. Thus, employing the ENCODE blacklists to filter C&R data might lead to both the undesired consequences of unnecessarily removing regions of true signal, and failing to remove C&R-specific artifacts.

**Figure 2:**
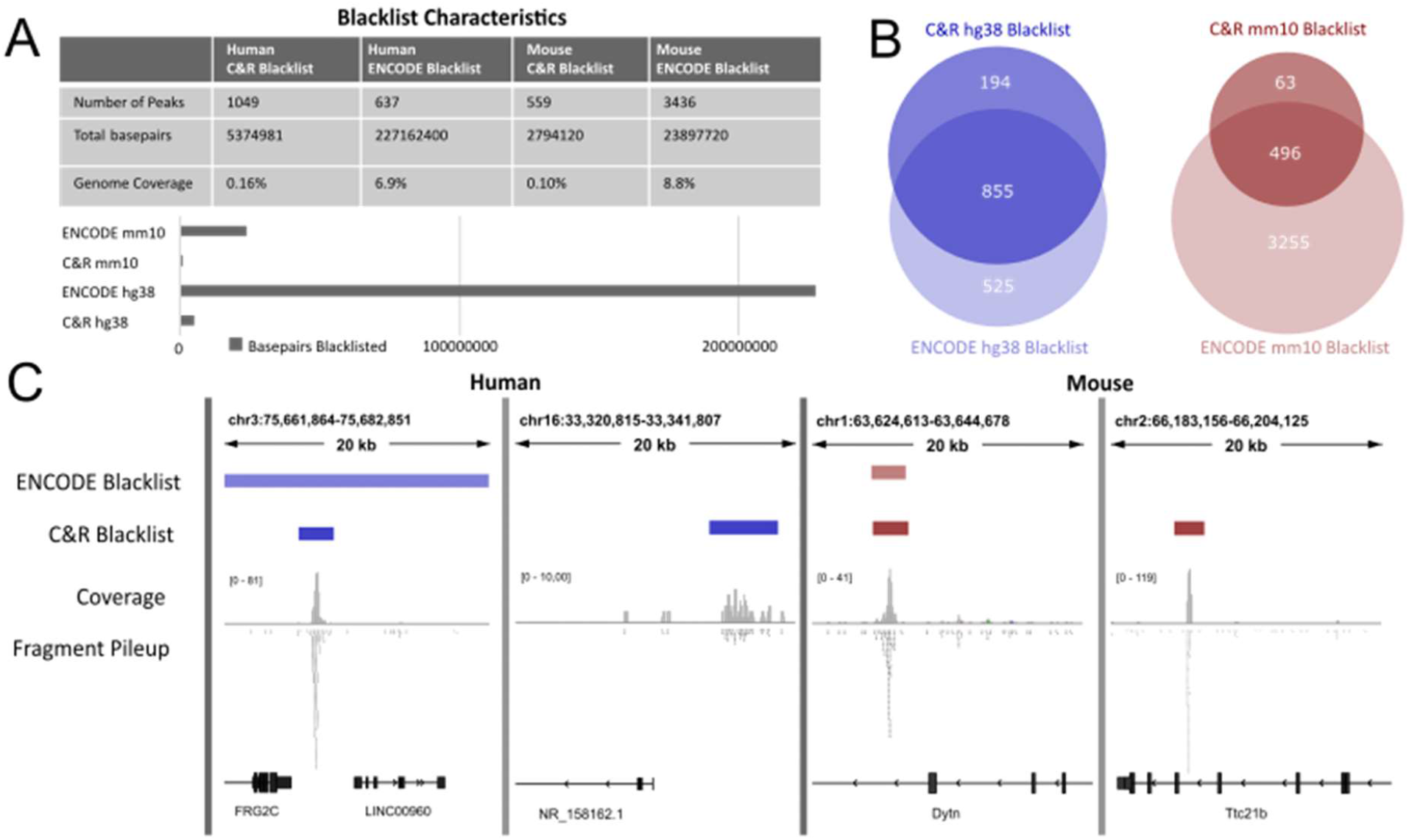
Comparison of the CUT&RUN and ENCODE Blacklists. **A.** Comparison of number of peaks, base pairs blacklisted, and total genome coverage for C&R and ENCODE hg38 and mm10 blacklists. The C&R blacklists cover less of the genome compared to their ENCODE counterparts. **B.** Comparison of overlapping blacklisted regions between C&R and ENCODE hg38 and mm10 blacklists, showing both considerable overlap and unique regions. **C.** Examples of coverage and fragment pileups for representative negative controls tracks in human and mouse, showing both unique regions for the C&R blacklists and regions shared with ENCODE. C&R = CUT&RUN

### Experimental Validation of the CUT&RUN Blacklists

To validate the C&R blacklists, we first turned to the negative controls they were built on. Principal component analysis (PCA) plots of the genome-wide signal distribution revealed a reduction in sample similarity after filtering with the blacklists, both for human and mouse C&R datasets (Figure 3A). This indicated that the C&R blacklists successfully identify and subtract the commonalities across negative control sample, which by definition should be considered false positives. In addition, to test our C&R blacklists experimentally, we decided to carry out a series of new experiments: we performed 8 C&R tests in HEK293T human cells and 8 in mouse embryonic tissues from JAX Swiss mice by using either IgG or anti-HA antibodies. To increase diversity in this test, we used both the original C&R protocol [1] and our recently developed C&R-LoV-U version for non-DNA-binding transcriptional co-factors [11]. A full list of sample information is provided in Materials and Methods Table 2. The obtained datasets were analyzed as shown in Figure 1A to replicate the construction of the blacklists but solely built on these new data. The C&R blacklists were able to increase the variance in our internally generated negative controls (Figure 3B). Most importantly, our new datasets contained > 80% of the regions identified by each C&R blacklist (Figure 3C), indicating that the C&R blacklists i) contain reproducibly enriched regions, ii) are generally applicable and iii) do not contain any obvious cell type-specific bias. In our opinion, the additional peaks of our internally generated series of negative controls not found by the blacklists are likely due to HEK293- or mouse strain-specific enrichment (Figure 3D). This reinforces the need for performing negative controls at each experiment, in addition to using our blacklists.

**Figure 3:**
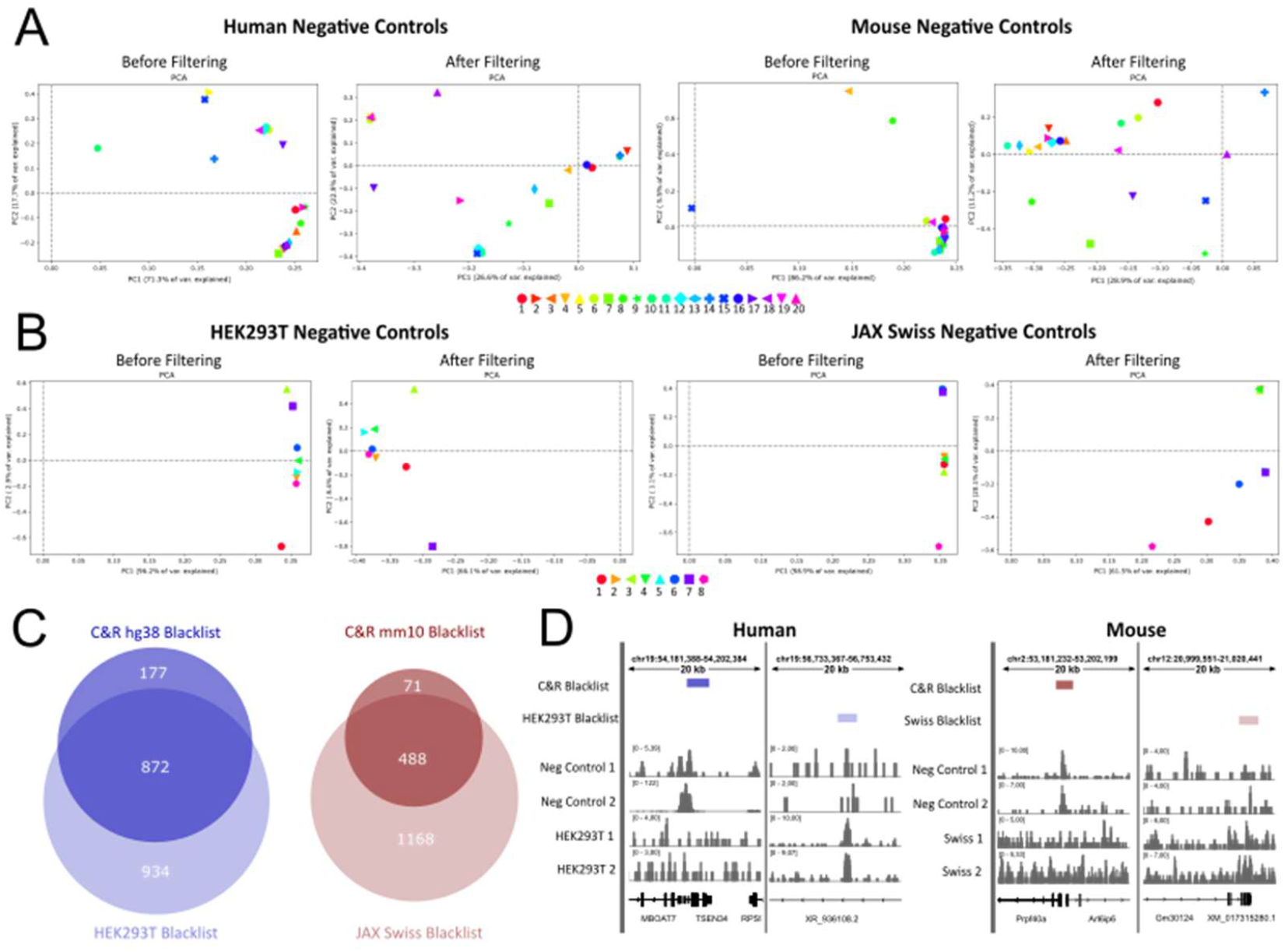
Experimental Validation of the CUT&RUN Blacklists. **A.** Principal component analysis graphs of human and mouse negative control samples, showing a decrease in sample similarity after filtering with C&R blacklists. The percentage of variance explained by each principal component decreases, and the samples become more randomly distributed on the graphs, indicating more variance in line with an expected random distribution of reads from a non-specific antibody in C&R. **B.** Principal component analysis on human and mouse negative controls performed to validate the C&R blacklists, showing increase variance after filtering. **C.** Comparison of C&R blacklists with blacklists built with the same method based on experiments performed in HEK293T human cells and embryonic tissues from JAX Swiss mice. The blacklists show high concordance with over 80% of each C&R blacklist being identified by the cell or strain specific blacklists. **D.** Examples of genome coverage of representative samples under unique regions of the C&R and cell/strain specific blacklists. C&R = CUT&RUN, PCA = Principal Component Analysis

**Table 2:** C&R Experiment Sample Information.

| Human |  |  |  | Mouse |  |  |  |
| --- | --- | --- | --- | --- | --- | --- | --- |
| Number | Condition | Protocol | Antibody | Number | Tissue | Protocol | Antibody |
| 1 | None | C&R | HA | 1 | Hindlimb | C&R LoV-U | IgG |
| 2 | Chir 4 h | C&R LoV-U | HA | 2 | Forelimb | C&R | IgG |
| 3 | Chir 24 h | C&R LoV-U | IgG | 3 | Branchial Arches | C&R | IgG |
| 4 | LGK | C&R | HA | 4 | Heart | C&R | IgG |
| 5 | Chir 24 h | C&R | HA | 5 | Hindlimb | C&R | IgG |
| 6 | Chir 24 h | C&R | IgG | 6 | Forelimb | C&R LoV-U | IgG |
| 7 | Chir 24 h | C&R LoV-U | IgG | 7 | Branchial Arches | C&R LoV-U | IgG |
| 8 | Chir 4 h | C&R LoV-U | IgG | 8 | Forelimb | C&R | HA |

### Filtering via CUT&RUN Blacklists Improves Peak Signal Intensity of Published Datasets

We set out to evaluate the performance of the C&R blacklist on published C&R datasets. First, we performed PCA analysis before and after blacklist filtering on human and mouse C&R-LoV-U datasets obtained across different types of targets, histone modifications, transcription factors, and co-factors [11]. This invariably led to similar improvements: first, the genome-wide variance of the signal increased; second, the samples clustered more, as expected, with their replicates or DNA binding partners. These indicate that the blacklist regions were falsely inflating the similarities of the samples (Figure 4A). We noticed that some of the blacklisted regions even appeared, in the published description of this experiment, within the list of “high-confidence” peaks for β-catenin in both human cells and mouse tissue, despite that these lists were curated by the subtraction of regions enriched in the negative control and of the ENCODE blacklists, enforcing the need for C&R specific blacklists.

**Figure 4:**
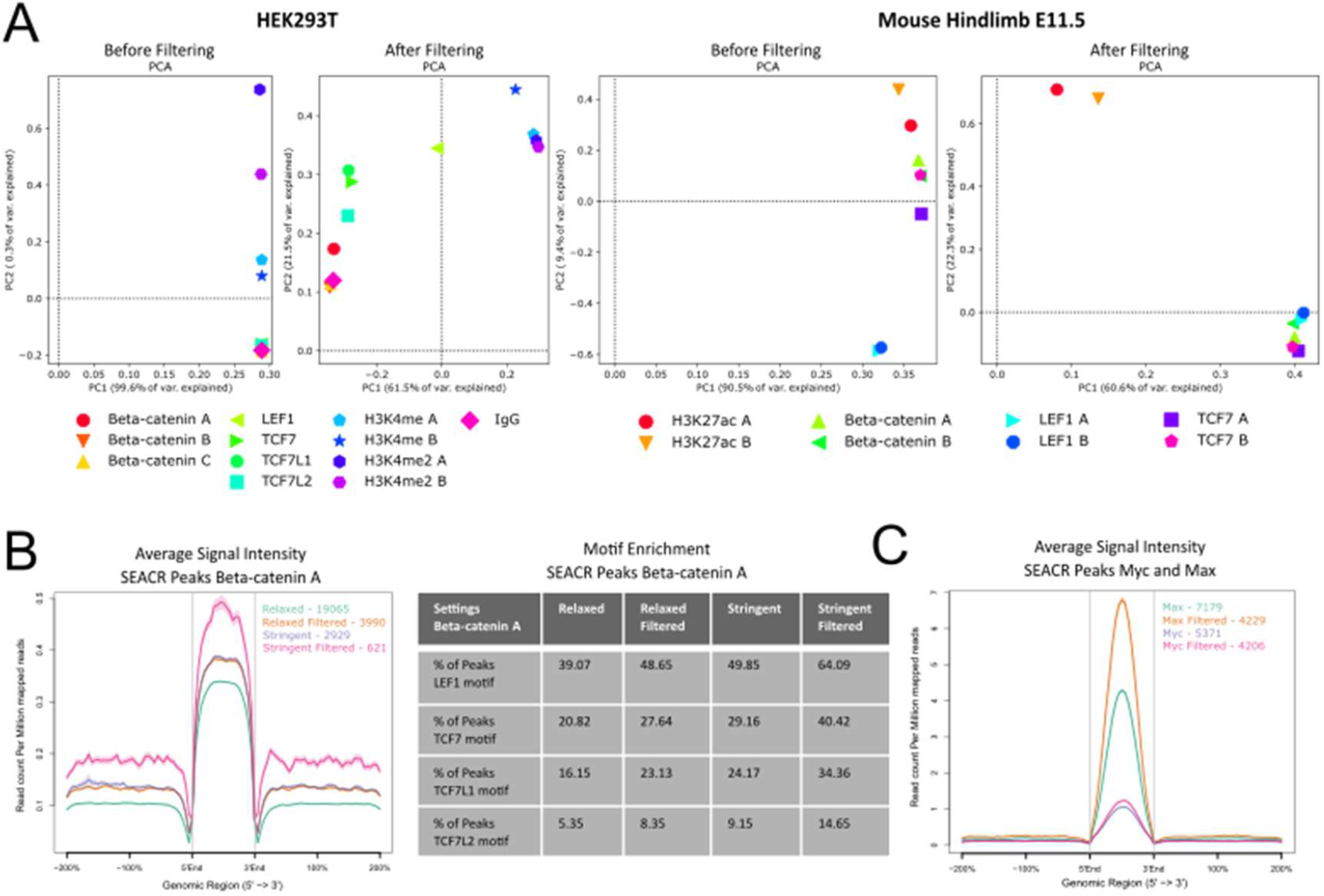
Application of the CUT&RUN Blacklists on Published Datasets. **A.** Principal component analysis on published C&R datasets from [11], showing that blacklist filtering reduces sample similarity and allows samples to clustering as expected based on antibody used. **B**. Comparison of peaks called using SEACR against a negative control on relaxed and stringent modes for the β-catenin A replicate, showing that blacklist filtering of BAM files before peak calling leads to a smaller number of called beaks which have an increased average signal profile within the peaks (left) and an increased percentage of peaks which contain expected TCF/LEF motifs (right). **C.** Comparison of peaks called by SEACR on Myc and Max datasets from [1] before and after blacklist filtering, showing a decrease in peak number accompanied by an increase in average signal within the peak regions. C&R = CUT&RUN, PCA = Principal Component Analysis

SEACR was benchmarked on datasets without removing high-signal regions, and often performs well as is on high-efficiency experiments. However, we hypothesized that signal within blacklist regions could be affecting the peak calling algorithm, both in the normalization of sample to control and in the later determination of signal thresholds. We tested SEACR on the β-catenin data with both relaxed and stringent settings and show that removing all the reads mapping to the C&R blacklisted regions before peak calling results in smaller, yet more reliable peak sets with fewer false positives. We conclude this based on the measurement of two key parameters: i) a higher enrichment for the expected motifs and ii) an improved average signal profile in the peaks (Figure 4B). Interestingly, analogous blacklist filtering had a relatively smaller effect on peak calling when this is performed with MACS2 on these data, aside from the identification of a number of blacklist regions as peaks when filtering was not first applied. This implies that SEACR, which is based on the detection of enrichment regions that stand out against the global background, is particularly sensitive to spurious signal.

To test this further, we downloaded the raw data for Max and Myc chromatin profiling performed in human K562 cells, along with the no-antibody negative control, from the original C&R publication [1]. First, we mapped these datasets with bowtie2 to the hg38 genome with and without deduplication, then calculated FRIP scores for the C&R hg38 blacklist. The Myc and no-antibody datasets displayed enrichment for the blacklist regions, with FRIP scores ranging from 5 to 8%. The Max dataset was especially affected by the blacklist regions: it had 71.9% of fragments within the blacklist peaks before deduplication, and 21.6% after duplicate removal. When called with SEACR against the negative control, before and after blacklist filtering, we detected a reduction in peak number for both Myc and Max, and an increased average signal-to-noise ratio within the peaks (Figure 4C). This difference is especially strong for Max, which had the higher FRIP score, and thus had likely a greater effect from signal within the blacklist regions.

When SEACR is performing well, it may not be needed to filter out blacklist regions. However, when SEACR calls very few or excessive numbers of peaks, we find that filtering can significantly improve this issue. We submit that removal of the C&R blacklist, whether before or after peak calling, is a useful tool to increase reliability of peak sets.

### Limitations

While the available published C&R data is increasing, there is no central repository like the ENCODE project for C&R data from which we could easily extract data to build a blacklist. In searching for data to use for the blacklist generation, we came across many datasets that either completely lacked negative controls or did not possess a clear sample labelling system, preventing us from their inclusion. We parsed through approximately 200 projects on the ENA before finding 20 available independent datasets each for human and mouse to use. Thus, our C&R blacklists were built on comparably less data than others previously published blacklists, and this might limit the robustness of the blacklists generated here. However, with the diversity of data used to build the blacklists and their subsequent successful validation, we believe we have compiled a mature selection of artifact regions that are a typical by-product of C&R experiments and are cell-type agnostic. We cannot conclusively exclude that our blacklists contain some regions that could be cell-type or strain specific or fail in identifying additional problematic regions that would appear in other cell types. Nevertheless, in the light of the many publicly available datasets lacking negative controls, we believe that our blacklist is an essential tool, as it improves the reliability of all C&R datasets tested. However, we also suggest that the blacklists generated here should accompany, rather than replace, adequate negative controls performed in the same conditions as the test sample.

## Methods

### Blacklist Generation

Publicly available C&R data: 40 negative controls (20 for human and 20 for mouse) were downloaded from the ENA archive, a full list of references and sample information is shown in Table 1. Read trimming was performed using bbmap bbduk [12] (version 38.18) removing adapters, artifacts, and poly AT, TA, G and C repeats. Alignment was done to the hg38 genome or mm10 genome using bowtie2 (version 2.4.5) [13], settings included --local --very-sensitive-local --no-unal --no-mixed --nodiscordant –dovetail --X 700. For bowtie [14] (version 1.0.0) we used the options -v 0 -m 1 -X 500. Samtools [15] (version 1.11) was used to create bam files, mark and remove duplicates when applicable, and sort bam files. Bedgraphs were created using bedtools [16] (version 2.23.0), genomecov with pair-end settings. Peaks were called using SEACR [8] (version 1.3) for each bedgraph using the settings norm and stringent with a threshold set to 0.001. Peak regions were extended using bedtools slop to 1000 bp in either direction from the peak. Bedsect [17] was used to overlap peak regions on default settings, and peaks called in greater than 30 % of negative controls (>= 7 of 20) were kept. Bedtools merge and sort was used to combine overlapping peaks to generate the final blacklist.

### Graphs and Analysis

deepTools [56] (version 3.5.1-0) multiBamSummary on bins mode for whole-genome, and bed mode using the blacklist bed file, followed by plotCorrelation were used to create correlation heatmaps with Spearman’s correlation. FRIP scores were calculated from total number of fragments mapping to blacklisted peaks, divided by total number of mapped fragments. Signal intensity plots and average profiles were created using ngsplot [57] (version 2.63) with the settings -G hg38 -R bed - N 2 -SC global -IN 0. The UCSC genome browser was used to create the chromosome graph [58]. Intervene [59] (version 0.6.4) was used to overlap peak sets and create Venn diagrams. IGV [60] was used for bedgraph and bed file visualization. The ENCODE v2 blacklists were downloaded from [6], and the mitochondrial genome was manually added for a fair comparison.

### Blacklist Validation on Published Data

Datasets were downloaded from [11] and [36]. Human and mouse data from Zambanini et al. [11], were processed as previously described in the paper. Raw data for Max, Myc and no antibody negative control in K562 cells from [1] were processed with bowtie2 as previously described, with or without removal of duplicates. Genome wide PCA plots were generated by first using deepTools multiBamSummary as described above on original data and by using -bl to filter out the C&R blacklists, and then the graphs made with plotPCA with default settings. Blacklist filtering prior to peak calling was performed on the BAM files with bedtools intersect -v. Peak calling was performed with SEACR against the negative control with the settings norm and relaxed or norm and stringent. Signal profile graphs were created with ngsplots with the settings -G hg38 -R bed -N 2 -SC global -IN 0. Motif analysis was done using Homer [61] (version 4.11) findMotifsGenome with the - size given option.

### CUT&RUN Experiments

Human negative control data was obtained from experiments on HEK293T cells. Cells were cultured at 37 °C in a humidified incubator with 5% CO2 in high glucose Dulbecco’s Modified Eagle Medium (Cat. #41965039, Gibco) supplemented with 10 % bovine calf serum (Cat. #1233C, Sigma-Aldrich) and 1X Penicillin-Streptomycin (Cat. #15140148, Gibco). Cells were stimulated with 10 μM CHIR99021 (Cat. #SML1046, Sigma Aldrich), 1 nM LGK (Cat. # S7143, Selleck Chemicals) or no stimulation. Mouse negative control data was obtained from various tissues from JAX Swiss Outbred mice (strain #: 034608) embryos at 11.5 dpc. Animal experimentation and housing conditions were according to the Swedish laws and guidelines under and performed under the ethical animal work license obtained by C.C. at Jordbruksverket (Dnr 2456-2019). Tissue dissociation, cell harvest, CUT&RUN or CUT&RUN LoV-U, and library preparation were performed as described in [11], CUT&RUN based on the original method by [1]. Anti-IgG (ABIN101961) or anti-HA (05-902R, Merck Millipore) antibodies were used. Samples were sequenced 36 bp pair-end on the NextSeq 550 (Illumina) using the Illumina NextSeq 500/550 High Output Kit v2.5 (75 cycles) (Cat. #20024906, Illumina). Sample information provided in Table 2.

## Supporting information

C&Rblacklist_hg38

C&Rblacklist_mm10

## Data Availability

Publicly available datasets used in this study can be downloaded according to their accession information in Table 1. C&R negative control experiments performed have been deposited at ArrayExpress, E-MTAB-12411. The C&R Blacklists are provided as supplementary files in .xls and .bed formats.

## Acknowledgements

This work was supported by Grants to C.C. from Cancerfonden (CAN 2018/542 and 21 1572 Pj), the Swedish Research Council, Vetenskapsrådet (2021-03075), and by Linköping University. C.C. is a Wallenberg Molecular Medicine (WCMM) fellow and receives generous financial support from the Knut and Alice Wallenberg Foundation. The computations and data handling were enabled by resources provided by the Swedish National Infrastructure for Computing (SNIC) at [SNIC CENTRE] partially funded by the Swedish Research Council through grant agreement no. 2018-05973.

## Contributions

AN identified the necessity for the tool, performed the experiments, carried out bioinformatics analyses, and wrote the manuscript and prepared the figures. GZ and PP performed the experiments and contributed with continuous feedbacks and discussions. CC supervised the research team, provided critical oversight of the analyses, and edited the manuscript. All authors contributed to multiple rounds of figures and manuscript revision.

## Notes

### Competing Interest Statement

The authors have declared no competing interest.

